# *Hnf1b*-CreER causes efficient recombination of a Rosa26-RFP reporter in duct and islet δ cells

**DOI:** 10.1101/2021.05.23.445318

**Authors:** Meritxell Rovira, Miguel Angel Maestro, Vanessa Grau, Jorge Ferrer

**Author notes:** Corresponding authors: Meritxell Rovira, Jorge Ferrer. Equal contribution (Meritxell Rovira, Miguel Angel Maestro).

## Abstract

The *Hnf1b*-CreER^T2^ BAC transgenic (Tg(Hnf1b-cre/ERT2)1Jfer) has been used extensively to trace the progeny of pancreatic ducts in developmental, regeneration, or cancer models. *Hnf1b*-CreER^T2^ transgenics have been used to show that cells from that form a duct-like plexus in the embryonic pancreas are bipotent duct-endocrine progenitors, whereas adult mouse duct cells are not a common source of β cells in various regenerative settings. The interpretation of such genetic lineage tracing studies is critically dependent on a correct understanding of the cell type specificity of recombinase activity with each reporter system. We have re-examined the performance of *Hnf1b*-CreER^T2^ with a Rosa26-RFP reporter transgene. This showed inducible recombination of up to 96% adult duct cells, a much higher efficiency than previously used reporter transgenes. Despite this high duct-cell excision, recombination in α and β cells remained very low, similar to previously used reporters. However, nearly half of somatostatin-expressing δ cells showed reporter activation, which was due to Cre expression in δ cells rather than to duct to δ cell conversions. The high recombination efficiency in duct cells indicates that the *Hnf1b*-CreER^T2^ model can be useful for both ductal fate mapping and genetic inactivation studies. The recombination in δ cells does not modify the interpretation of studies that failed to show duct conversions to other cell types, but needs to be considered if this model is used in studies that aim to modify the plasticity of pancreatic duct cells.

## Main text

Understanding the lineage relationships of adult cell types has important implications in defining regenerative strategies for human diseases. If adult pancreatic progenitors or differentiated cells have the capacity to generate new insulin-producing β cells, this could be harnessed to stimulate β cell formation in type 1 diabetes. Likewise, insights into the cellular origin of different types of pancreatic cancer has major implications to understand and target early oncogenic processes.

Genetic lineage tracing can provide unequivocal evidence for lineage relationships. This can be carried out with a mouse allele that achieves specific expression of a Cre recombinase in a candidate progenitor cell type (Cre driver), combined with a second allele that undergoes Cre-based excision of a STOP cassette flanked by LoxP sites. Cre-mediated activation of the reporter (Cre responder) in the putative progenitor allows tracing all of its cellular progeny.

In several instances, genetic lineage tracing has challenged pre-existing conceptions that are rooted on the assumption that whenever a cell co-expresses markers from two different lineages it must be transitioning between both lineages, or represent some form of progenitor that can differentiate to different lineages. For example, the co-expression of insulin and glucagon in embryonic pancreatic cells was initially thought to indicate that β cells originate from common bi-hormonal cells, but lineage tracing studies later disproved this notion, showing that insulin and glucagon-expressing cells arise as separate lineages. ^1^ Likewise, insulin-expressing cells that reside in the adult pancreatic ductal epithelium and co-express cytokeratin have often been referred to as duct-derived neogenic β cells. Such a putative postnatal source of adult β cells aligns well with the fact that duct-like cells from the embryonic plexus are bipotent progenitors of endocrine and duct cells. ^2-5^ The hypothesis that adult duct cells are facultative progenitors was tested using a BAC transgenic that contains ∼190 Kb of the *Hnf1b* locus and drives an inducible Cre recombinase in pancreatic duct cells (*Hnf1b*-CreER^T2^, or Tg(Hnf1b-cre/ERT2)1Jfer). ^2^ These experiments, which used a Rosa26-β galactosidase (βgal) reporter, concluded that –contrary to earlier beliefs– mouse adult pancreatic duct cells are not a common source of β cells during the normal lifespan of mice, or under several regenerative settings that have been proposed to result in duct to endocrine transdifferentiation. ^2^ Similar results were obtained with other duct Cre lines– including *Hes1* ^6^, *Sox9* ^3^ and *Mucin1* ^7^, and are compatible with β cell or dual β cell/non β cell lineage tracing in regeneration models.^8;9^ Thus, with the exception of studies using the CAII promoter ^10^, genetic lineage studies have suggested that duct-like cells give rise to endocrine cells throughout development but are not a common pool of endocrine progenitors in the adult pancreas. It is nevertheless possible that pancreatic duct cells can be manipulated to transdifferentiate into β cells as a replacement therapy for diabetes.

Genetic lineage tracing is critically dependent on careful design and precise knowledge of which cell types are labeled at the outset of the labeling experiment. Failure to define this parameter can lead to erroneous conclusions concerning the cellular origins of cells being examined. Incomplete knowledge of cell types marked by various Cre models could, for example, have contributed to some discordances with different duct Cre tracer models. The *Hnf1b*-CreER^T2^ BAC transgenic line, in particular, has been used by numerous publications to study the progeny of HNF1B-expressing cells in a wide range of experimental models, including cancer, dynamics of cellular differentiation, or regeneration. ^11-14^ This underscores a need for a more exhaustive analysis of its performance.

In the current study we have extended the characterization of *Hnf1b*-CreER^T2^ BAC transgenic model using Rosa26-RFP instead of two previously employed reporter lines, Rosa26YFP and Rosa26-βgal. With the Rosa26-RFP reporter, Cre-based excision of a stop cassette leads to expression of a concatemerized red fluorescent protein (RFP) that displays high brightness and can thus be used for live imaging. ^15^

*Hnf1b*-CreER^T2^ - Rosa26-βgal 8-12 week old mice were treated with tamoxifen (three doses of tamoxifen –20 mg, 20 mg, and 10 mg– by gavage every other day), and analyzed one week later typically showed ∼20% recombination efficiency in Cytokeratin 19-labeled pancreatic duct cells (Solar et al.^2^ and **Figure 1A**). The same treatment in *Hnf1b*-CreER^T2^ - Rosa26-RFP mice led to 55-95% recombination efficiency in duct cells (73.6 ± 6.3) (**Figure 1B and E**). Thus, the Rosa26-RFP reporter enabled much higher recombination efficiency in adult duct cells than previously reported with the same Cre allele. We believe that this was likely caused by differences in the recombination efficiency of the Rosa26-RFP reporter, rather than by differences in the sensitivity to detect the RFP reporter, because duct cell labeling with Rosa26-RFP was also substantially higher than with a Rosa26-YFP reporter system that uses the similarly sensitive immunodetection method^2^.

**Figure 1:**
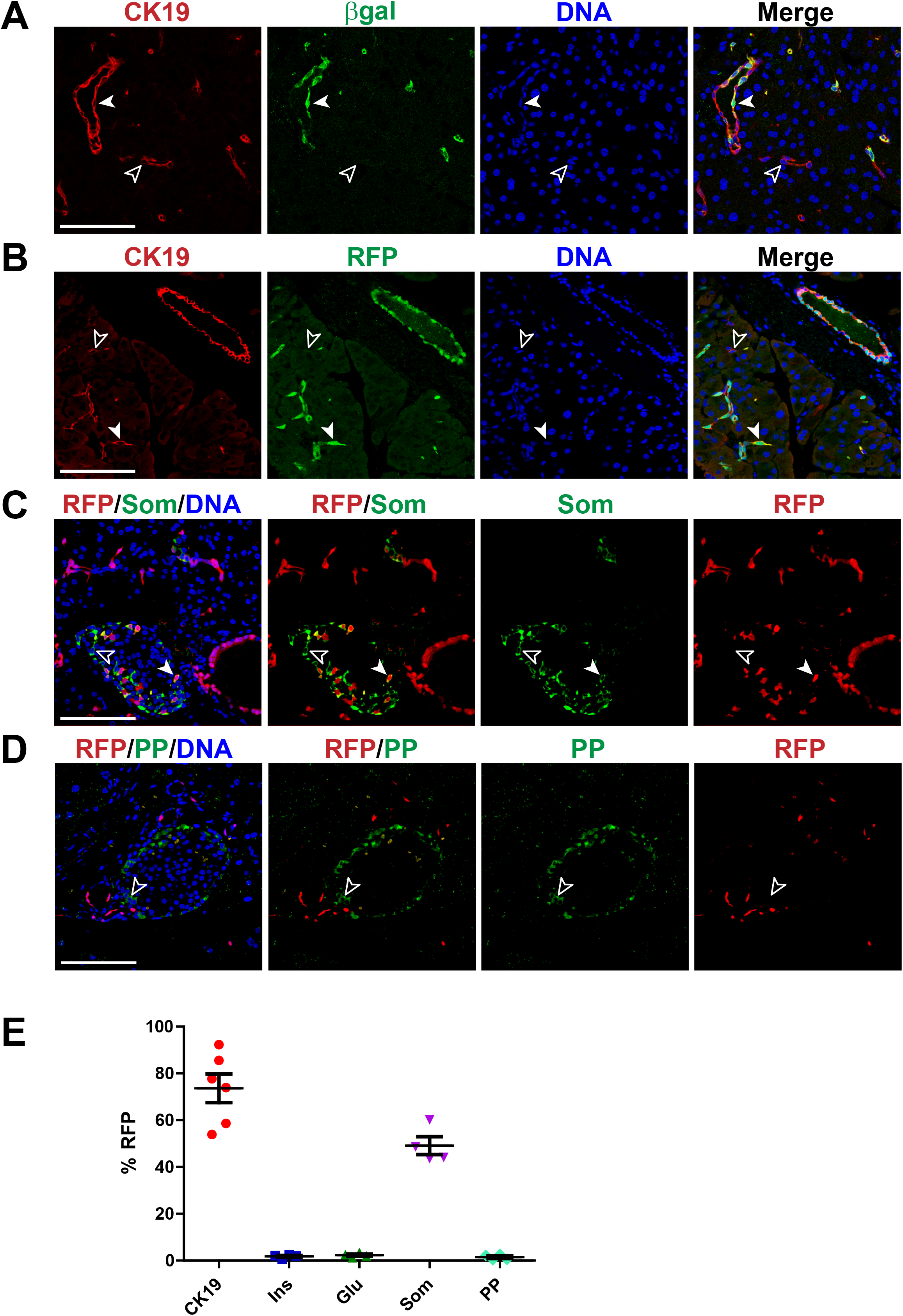
Efficient recombination in pancreatic ducts and labeling specificity in endocrine lineages. (A) Immunofluorescence for β-galactosidase (green), Cytokeratin 19 (CK19, red) and DAPI (blue) in adult mouse pancreas shows moderate efficiency of recombination of the *Hnf1b*-CreER^T2^;Rosa26-β-gal transgenics in duct cells. (B) Immunofluorescence for RFP (green), Cytokeratin 19 (CK19, red) and DAPI (blue) in adult mouse pancreas shows high efficiency of recombination of the *Hnf1b*-CreER^T2^;Rosa26-RFP transgenics in duct cells. (C) Immunofluorescence for RFP (red), somatostatin (Som, green) and DAPI (blue) in adult mouse pancreas shows recombination in somatostatin positive δ cells in *Hnf1b*-CreER^T2^;Rosa26-RFP transgenics. (D) Immunofluorescence for RFP (red), pancreatic polypeptide (PP, green) and DAPI (blue) in adult mouse pancreas shows very low recombination of PP-expressing cells in *Hnf1b*-CreER^T2^;Rosa26-RFP transgenics. White arrowheads indicate double positive cells while empty arrow heads indicate single positive cells. (E) Percentage of RFP positive cells in duct (n=6 mice), β (n= 4 mice), α (n= 4 mice), δ (n= 4 mice) and PP (n= 4 mice) positive cells. Antibodies used in the study: RFP (Rockland #600-401-379S, 1:500), CK19 (Hybridoma Bank #Troma-III, 1:50), Insulin (Linco #4011-01F, 1:1000), Glucagon (Sigma #4031-01F, 1:1000), Somatostatin (Santa Cruz #sc-7819, 1:50), PP (Novus #NB100-1793, 1:200). Scale Bar=100 μm.

This increased recombination efficiency when using a different reporter line led us to re-examine the pancreatic cell type specificity of *Hnf1b*-CreER^T2^-Rosa26-RFP mice with the same tamoxifen treatment. We found that β and α cell labeling with Rosa26-RFP was not substantially higher than previously reported; 1.82% ± 0.44 of β cells were RFP+ and 2.28% ± 0.5 of α cells were RFP+ (**Figure 1E)**. For comparison, with Rosa26-βgal we found that 1.04% ± 0.48 of β cells were βgal+, and 1.22% ± 0.69 of α cells were βgal+, which was similar to 1-2% labeling rates previously reported with *Hnf1b*-CreER^T2^-Rosa26-βgal. ^2^ We observed a similar very low recombination of insulin+ and glucagon+ cells amongst rare hormone producing cells that line the ductal epithelium. Thus, despite the marked increase in labeling efficiency of adult ducts with the new reporter, there was no increase in the number of α and β cells carrying the label.

Our original studies focused on labeling of β and α cells because they are by far the most prevalent islet endocrine cells, and because those studies needed to ensure if the model was valid for testing the hypothesis that duct cells give rise to β cells. Analysis of tamoxifen-treated *Hnf1b*-CreER^T2^-Rosa26-RFP mice, however, revealed conspicuous RFP+ cells in the mantle of some pancreatic islets, and were glucagon/insulin-negative, which led us to examine other hormones. Somatostatin-expressing δ cells make up 1-5% of islet endocrine cells.^16^ We found that on average 49.5% ± 4.1 of δ cells were RFP+ in *Hnf1b*-CreER^T2^;Rosa26-RFP islets (see **Figure 1C**, showing an islet with multiple δ cells, and quantifications in **Figure 1E)**. By contrast, only 1.45% ± 0.6 of pancreatic polypeptide cells were RFP+ (**Figure 1D** and **E**).

Labeling of δ cells by the *Hnf1b*-CreER^T2^ model could mean that Cre recombination took place in δ cells at the outset of tamoxifen treatment because CreER is expressed in these cells, or else that recombination occurred in HNF1B-expressing duct cells which later gave rise to δ cells. We thus analyzed Cre expression after a short (24 hour) tamoxifen treatment pulse. Tamoxifen induces nuclear translocation of Cre. We observed widespread nuclear Cre protein expression in cells that line the duct epithelium (**Figure 2A**) as well as unequivocal expression of Cre protein in δ cells, but not in other islet cells (**Figure 2B**). Because *Hnf1b*-CreER^T2^ is a large ∼190 Kb BAC transgenic model that is expected to recapitulate endogenous expression of *Hnf1b*, we next examined if Cre expression reflected the expression pattern of the endogenous HNF1B protein in δ cells. HNF1B immunostaining was not clearly detected in endocrine cells including δ cells, in sharp contrast with HNF1B-expressing duct cells (**Figure 2C**).

**Figure 2:**
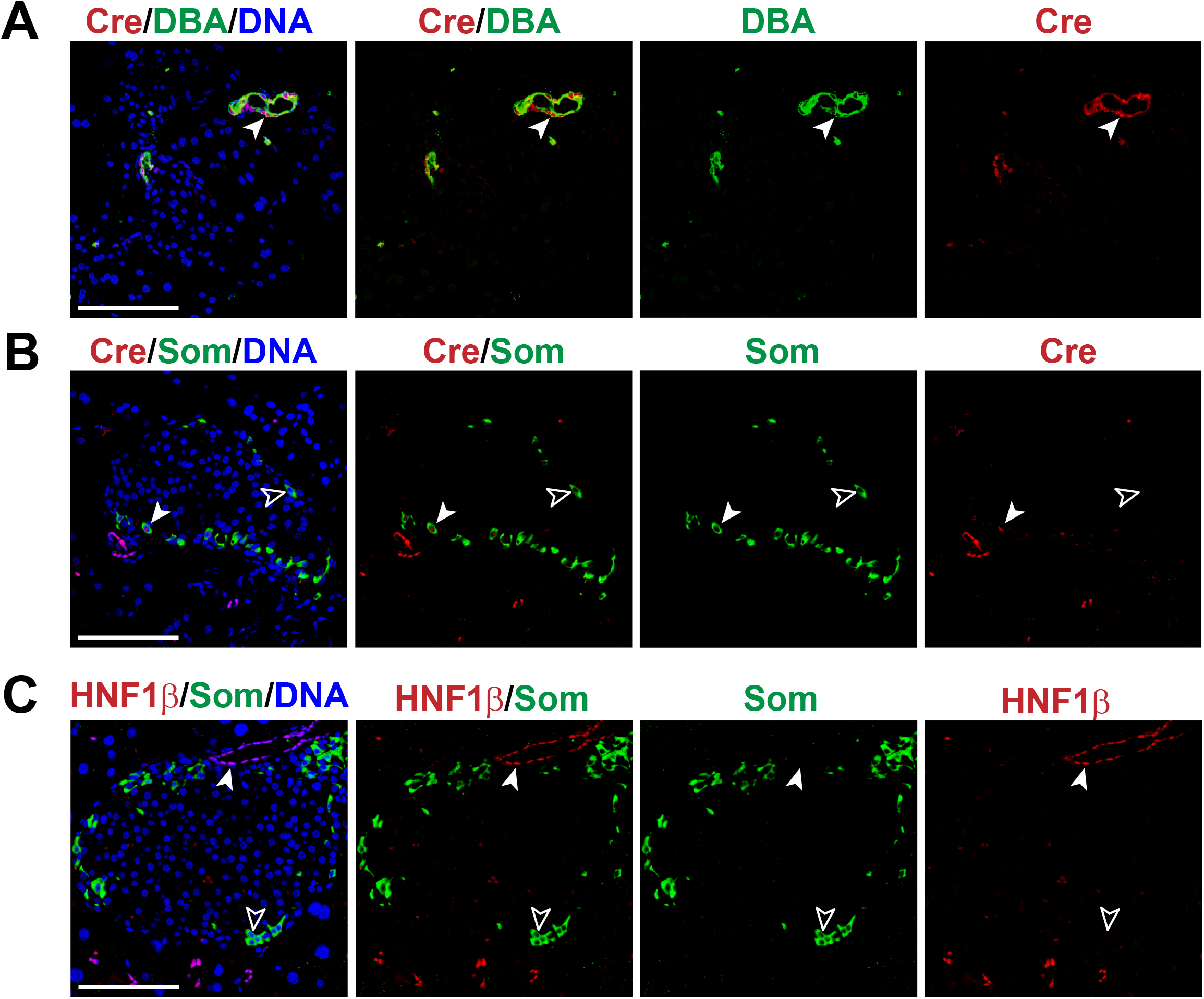
Cre is expressed in δ cells. (A) Immunofluorescence for Cre (red), Dolichos Biflorus Agglutinin (DBA, green) and DAPI (blue) in adult mouse *Hnf1b*-CreER^T2^;Rosa26-RFP pancreas upon 24 hours of tamoxifen treatment shows high Cre expression in duct cells. (B) Immunofluorescence for Cre (red), somatostatin (Som, green) and DAPI (blue) in adult transgenic mouse pancreas upon 24 hours of tamoxifen treatment shows Cre expression in several δ cells. (C) Immunofluorescence for HNF1B (red), somatostatin (Som, green) and DAPI (blue) in adult transgenic mouse pancreas upon 24 hours of tamoxifen treatment shows strong expression of HNF1B in ductal cells but not in somatostatin positive cells. White arrowheads indicate doble positive cells while empty arrow heads indicate single positive cells. Antibodies used in the study: DBA (vector laboratories #FL-1031, 1:200), Hnf1b (Santa Cruz #sc-22840, 1:200), Cre (Novagen #69050-3, 1:1000), Somatostatin (Santa Cruz #sc-7819, 1:50). Scale bar= 100 µm.

These findings, therefore, indicate that *Hnf1b*-CreER^T2^ marks adult pancreatic δ cells, and this can be ascribed to expression of Cre in these cells. The expression of Cre, but not HNF1B, could be due to higher transcriptional activity of *Hnf1b* in δ cells that is not reflected by immunofluorescence due to cell-specific post-transcriptional regulation of protein levels, or to cell-specific cis-acting silencing sequences that are not included in the 190 Kb transgenic BAC.

The high labeling efficiency in δ cells contrasts with a very low recombination frequency and lack of visible Cre expression in α and β cells. Of note, the rare labeling of α and β cells is also ascribed to direct recombination in these cells because it is present immediately after Tamoxifen treatment and does not change over the course of months or experimental perturbations^2^. Plausibly, α and β recombination obeys to very weak transcriptional activity of the *Hnf1b* fragment in α and β cells, leading to Cre expression that is too low for immunodetection but can cause occasional excision of reporter alleles. This background recombination rate needs to be monitored in studies that use this model.

The new findings reported here should be considered to interpret lineage tracing studies of adult duct cells and their progenitor potential. For studies that have not observed duct to β cell transitions in various settings, this does not influence conclusions, except that they extend the intepretation in that neither duct cells nor any δ cells that might have been labelled in adult mice give rise to other cells examined under those conditions. However, studies that aim to elicit cellular conversions from ductal epithelium and use this model need to consider this alternate source of excised cells.

The results also show higher ductal excision rates than previously reported with the same Cre line. The efficiency of Cre-based excision can vary considerably for different LoxP alleles, which is often influenced by easier excision of shorter LoxP-flanked fragments. Our results indicate that the *Hnf1b*-CreER^T2^ model can be used to ablate genes in the pancreatic duct epithelium provided that the target locus allows for high recombination, keeping in mind that δ cells are also expected to recombine. The findings also indicate that Rosa26-RFP reporter can be used when higher efficiency of duct cell labeling is desired.

Finally, it is interesting to note that the widespread, sometimes nearly exhaustive, ductal excision did not increase the proportion of labeled β cells, including those that reside in ducts, showing that the ductal location of an islet cell should not be taken as evidence of a recent duct to β cell conversion.

## Acknowledgements

This research was supported by from Ministerio de Ciencia e Innovación (MCIU) (SAF2015-73226-JIN and RYC-2017-21950) and (FP-PEOPLE-2007-2-3-COFUND) to M.R. and grants to J.F. from Ministerio de Ciencia e Innovación (BFU2014-54284-R, RTI2018-095666-B-I00). Work in CRG was supported by the CERCA Programme, Generalitat de Catalunya and Centro de Excelencia Severo Ochoa (SEV-2015-0510).

## Author Contributions

M.R., M.A.M., and J.F. conceived and coordinated the study. M.R., M.A.M. performed image analysis. V.G. maintained mouse colonies and performed tamoxifen treatments. M.R. and J.F. wrote the manuscript with input from the remaining authors.

